# AT-527 is a potent *in vitro* replication inhibitor of SARS-CoV-2, the virus responsible for the COVID-19 pandemic

**DOI:** 10.1101/2020.08.11.242834

**Authors:** Steven S. Good, Jonna Westover, Kie Hoon Jung, Paolo La Colla, Gabriella Collu, Adel Moussa, Bruno Canard, Jean-Pierre Sommadossi

**Author notes:** Corresponding Author: Steven S. Good.

## Abstract

AT-527, an orally administered double prodrug of a guanosine nucleotide analog, has been shown previously to be highly efficacious and well tolerated in HCV-infected subjects. Herein we report the potent *in vitro* activity of AT-511, the free base form of AT-527, against several coronaviruses, including SARS-CoV-2, the causative agent of COVID-19. In normal human airway epithelial (HAE) cell preparations, the average concentration of AT-511 required to inhibit replication of SARS-CoV-2 by 90% (EC_90_) was 0.5 µM, very similar to the EC_90_ for AT-511 against HCoV-229E, HCoV-OC43 and SARS-CoV in Huh-7 cells. No cytotoxicity was observed for AT-511 in any of the antiviral assays up to the highest concentration tested (100 µM). Surprisingly, AT-511 was 30-fold less active against MERS-CoV. This differential activity may provide a clue to the apparent unique mechanism of action of the guanosine triphosphate analog formed from AT-527.

## INTRODUCTION

A novel coronavirus, severe acute respiratory syndrome coronavirus-2 (SARS-CoV-2), was recently identified as the pathogen that causes the potentially life-threatening disease called coronavirus disease 2019 (COVID-19), which has rapidly spread across the world. Currently there are no vaccines or therapeutics approved to prevent SARS-CoV-2 infection or to treat COVID-19.

Recently remdesivir, the prodrug of an adenosine nucleotide analog, originally developed for the treatment of Ebola virus infections, has demonstrated potent *in vitro* activity against SARS-CoV-2 (Wang et al., 2020) and has shown sufficient promise in the treatment of COVID-19 patients (Beigel et al., 2020) such that the FDA has granted Emergency Use Authorization to this nucleotide prodrug. The limited oral bioavailability of remdesivir (Eastman et al., 2020), however, requires that it be administered via intravenous infusion, thus limiting its use to hospitalized patients. COVID-19 is an acute viral infection for which antiviral therapeutics will be most effective within the first stage of the infection when viral load is at its maximum, with rapid viral replication initially in nasal, throat and pulmonary cells (Mason, 2020). Only the availability of a potent, safe, oral antiviral administered to SARS-CoV-2 infected individuals in their early stage of disease will avert clinical illness and mitigate the COVID-19 pandemic.

AT-527, an oral double prodrug of a guanosine nucleotide analog, has demonstrated potent *in vitro* activity against clinical isolates of hepatitis C virus (HCV) by inhibiting the RNA-dependent RNA polymerase (RdRp) (Good et al., 2020). Supported by a comprehensive preclinical toxicology and safety pharmacology package, AT-527 has recently been evaluated in a Phase 1B study (Berliba et al., 2019) and a Phase 2 clinical trial (Mungar, et al. 2020). In the latter study, AT-527 was safe and well-tolerated for up to 12 weeks in HCV-infected subjects and achieved a high rate of efficacy. Amid the SARS-CoV-2 pandemic, the activity of AT-511, the free base of AT-527, was tested *in vitro* against several human coronaviruses, including SARS-CoV-2.

## RESULTS

In an initial screening, BHK-21 cells acutely infected with a seasonal human alpha coronavirus, HCoV-229E, were exposed to serial dilutions of AT-511. After a 3-day incubation, the effective concentration of AT-511 required to achieve 50% inhibition (EC_50_) of the virus-induced cytopathic effect (CPE) from two independent experiments averaged 1.8 µM (Table 1). In contrasst, the 2’-fluoro-2’-methyl uridine nucleotide prodrug sofosbuvir did not inhibit HCoV-229E replication at concentrations as high as 100 µM. No toxicity was detected from either drug.

**Table 1.** *In Vitro* Activity of AT-511 and Other Oral Antiviral Drugs Against Various Human Coronaviruses.

| Virus (genus) | Cell line | Compound | Neutral Red Assay |  | Virus Yield Reduction Assay<br>EC<sub>90</sub> (μM) | Selectivity Index<br>(CC<sub>50</sub>/EC<sub>90</sub>) |
| --- | --- | --- | --- | --- | --- | --- |
|  |  |  | EC<sub>50</sub> (μM) | CC<sub>50</sub> (μM) |  |  |
| HCoV-229E (alpha) | BHK-21 | AT-511 | 3.3<sup>a,b</sup> | >100<sup>b</sup> |  | >30 |
|  |  | sofosbuvir | >100<sup>b</sup> | >100<sup>b</sup> |  | N/A |
|  | Huh-7 | AT-511 | 1.7 / 1.6 | >86 | 1.0 | >86 |
|  |  | chloroquine | 8.1 | 21 | <0.050 | 2.6<sup>c</sup> |
|  |  | hydroxychloroquine | 7.4 | 26 | <0.048 | 3.5<sup>c</sup> |
| HCoV-OC43 (beta) | Huh-7 | AT-511 | ND<sup>d</sup> | >86 | 0.5 / <0.03 | >170 / >3100 |
|  | RD | AT-511 | 2.8 | >86 | 2.2 | >39 |
| MERS-CoV (beta) | Huh-7 | AT-511 | 15 / 36 | >86 | 17 / 56 | >5 / >1.5 |
| SARS-CoV (beta) | Huh-7 | AT-511 | ND | >86 | 0.34 | >250 |
| SARS-CoV-2 (beta) | HAE | AT-511 | ND | >86<sup>e</sup> / >8.6<sup>e</sup> | 0.64<sup>f</sup> / 0.47<sup>g</sup> | >130 / >18 |
|  |  | N<sup>4</sup>-hydroxycytidine |  | >19<sup>e</sup> | 3.9<sup>h</sup> | >4.8 |

<sup>a</sup>EC<sub>90</sub> (average of 2 experiments)
<sup>b</sup>Determined by MTT cell viability test
<sup>c</sup>CC<sub>50</sub>/EC<sub>50</sub> (virus yield reduction assay overestimates antiviral potency of cytotoxic compounds)
<sup>d</sup>Not determined (no cytopathic effect with this virus in this cell line)
<sup>e</sup>Cytotoxicity assessed by visual inspection of cell monolayers
<sup>f</sup>Average of two replicates (0.57 and 0.70 μM)
<sup>g</sup>Average of two replicates (0.52 and 0.42 μM)
<sup>h</sup>Average of two replicates (4.7 and 3.1 μM)
BHK-21, baby hamster kidney cell line
Huh-7, human hepatocyte carcinoma cell line (established ability to form triphosphate from AT-511)
RD, human rhabdomyosarcoma cell line (unknown ability to form triphosphate from AT-511)
HAE, normal human airway epithelial cell preparation (established ability to form triphosphate from AT-511)

The *in vitro* potency of AT-511 against HCoV-229E, HCoV-OC43 (another seasonal human coronavirus strain), MERS-CoV and SARS-CoV was then evaluated in Huh-7 cell-based assays. This human hepatocarcinoma cell line was selected based on its ability to activate AT-511 intracellularly to its triphosphate metabolite, unlike MRC-5 cells in which AT-511 lacked activity against HCoV-229E (EC_50_ >100 µM; Good et al., 2020). Antiviral activity was assessed by two different methods after exposure of Huh-7 cells to virus and serial dilutions of test compound by determining 1) the EC_50_ for virus-induced CPE by neutral red dye staining after a 5-day (229E and OC43) or 7-day (MERS and SARS) incubation and 2) the effective concentration required to reduce secretion of infectious virus into the culture medium by 90% (EC_90_) after a 3-day incubation using a standard endpoint dilution CCID_50_ assay to determine virus yield reduction (VYR). Half-maximal cytotoxicity (CC_50_) was measured by neutral red staining of compound-treated duplicates in the absence of virus. Although a robust VYR endpoint was obtained in Huh-7 cells infected with HCoV-OC43 or SARS-CoV, CPE was not observed and EC_50_ values using neutral red staining were not obtained with these viruses. Individual determinations of EC_90_ values for AT-511 against HCoV-229E, HCoV-OC43 and SARS-CoV ranged from 0.34 to 1.2 µM, whereas the value against MERS-CoV averaged 37 µM (Table 1). No cytotoxicity was detected with AT-511 up to 86 µM, the highest concentration tested.

Chloroquine and hydroxychloroquine appeared to be quite potent against HCoV-229E and HCoV-OC43 based on their EC_90_ values of <0.05 µM obtained using VYR measurements (Table 1). The respective EC_50_ values for these two drugs (8.1 and 7.4 µM), obtained using the neutral red assay, were substantially higher and only 2.6- to 3.6-fold less than the corresponding CC_50_ values, indicating considerably lower potencies and poor selectivity indices. These differences illustrate an inherent error in assessing antiviral activities of cytotoxic compounds using only measurements of VYR. When cells are poisoned by toxic drugs and are progressing toward death, their ability to support viral replication and propagation in addition to their own health likely is greatly diminished. At the point when cell death is detected by staining, viral yield reduction measurements likely reflect a combination of antiviral activity and cytotoxicity, thus overestimating antiviral potencies.

In contrast to a recent publication (Wang et al., 2020), Huh-7 cells were not permissive for replication of SARS-CoV-2. An assay was developed using normal human airway epithelial (HAE) cells, a highly relevant *in vitro* model of the lung, which has been established as a more representative system than cell lines for SARS-CoV-2 replication (Jonsdottir and Dijkman, 2016). These primary cells form polarized monolayers, the apical side of which is exposed to air and produces a mucin layer, consistent with the physiology of the human airways (Sheahan et al., 2020). Average EC_90_ and CC_50_ values for AT-511 against SARS-CoV-2 from two separate HAE assays (0.5 and >86 µM, respectively) were in the same range as those obtained for HCoV-OC43 and SARS-CoV (Table 1).

In the second HAE assay, the activity of AT-511 was tested in parallel with N^4^-hydroxycytidine (the nucleoside formed from the mutagenic oral prodrug EIDD-2801) with recently reported *in vitro* and *in vivo* activity against SARS-CoV-2 (Sheahan et al., 2020). The potency of N^4^-hydroxycytidine against SARS-CoV-2 (EC_90_ = 3.9 µM) was 8 times less than that of AT-511 in the same experiment.

## DISCUSSION

The 30-fold difference of AT-511 activity between MERS-CoV and other CoVs is puzzling. Nucleotide and nucleotide analogue selection is achieved at the CoV RdRp active site, the nsp12 gene product activated by its processivity co-factors nsp7 and nsp8 (Subissi et al., 2014). Conserved amino acid motifs A and C are involved in phosphodiester bond formation, whereas motifs F and B participate in nucleotide channeling and binding at the active site, respectively. No significant structural differences are apparent between MERS-CoV and other CoVs in these essential motifs. With a similar ribose modification between AT-511 and sofosbuvir, it is unlikely that the selective lack of activity of sofosbuvir would be due to excision by the CoV exonuclease carried by nsp14 (Ferron et al., 2018). Rather, our results suggest that the triphosphate formed from AT-511 most likely targets another less conserved viral GTP-binding protein, whose inhibition would account for both the antiviral effect and the MERS-CoV differential sensitivity pattern.

To date, human safety and tolerability have been confirmed in more than 40 HCV-infected patients treated orally with once-daily administration of 550 mg of AT-527 for seven days and up to 12 weeks. Excellent pharmacokinetics (50-60% bioavailability and long intracellular half-life of the active metabolite) associated with a dosing regimen of a 1,100 mg loading dose, followed by 550 mg BID, should provide lung drug exposures consistently above its *in vitro* EC_90_ of 0.5 µM against SARS-CoV-2 replication and should lead to a very effective antiviral treatment. A Phase 2 clinical trial is currently ongoing to evaluate the safety and efficacy of AT-527 in COVID-19 patients. Additionally, the differential activities AT-511 against MERS-CoV in comparison to the other CoVs tested and of AT-511 in comparison to sofosbuvir against HCoV-229E are being exploited to probe the unique mechanism of inhibition of the active guanosine triphosphate analog formed from AT-527.

## AUTHOR CONTRIBUTIONS

J.W. and K.J. developed and conducted the assays for HCoV-229E, HCoV-OC34, MERS-CoV and SARS-CoV-1 in Huh-7 cells and SARS-CoV-2 in HAE cell preparations. P.L.C. and G.C. developed and conducted the assays for HCoV-229E in BHK-21 cells. A.M. acquired the antivirals. S.G., A.M and J.S. designed the experiments. S.G., B.C. and J.S. wrote the manuscript. All authors provided input and final approval of the manuscript.

## MATERIALS AND METHODS

### Cells, viruses and compounds

BHK-21 (baby hamster kidney) cells, Huh-7 (human hepatocarcinoma) cells, RD (human rhabdomyosarcoma) cells and the seasonal human coronaviruses (HCoV-229E and HCoV-OC43) were obtained from American Type Culture Collection, Manassas, VA. MERS-CoV (EMC), SARS-CoV (Urbani) and SARS-CoV-2 (USA-WA1/2020) were supplied by The Centers for Disease Control and Prevention, Atlanta, GA. The HAE cell preparations (EpiAirway™ AIR-100 or AIR-112) were purchased from MatTek Corporation, Ashland, MA. AT-511 and N^4^-hydroxycytidine were prepared for Atea Pharmaceuticals by Topharman Shanghai Co., Ltd., Shanghai, China and Oxeltis, Montpellier, France, respectively. Chloroquine and hydroxychloroquine were purchased from Mason-Chem, Palo Alto, CA and sofosbuvir was purchased from Pharma Sys, Inc., Cary, NC.

### Antiviral assays in BHK-21 cells

Test compounds were dissolved in DMSO at 100 mM and then diluted in Minimum Essential Medium with Earle’s salts (MEM-E) containing 1 mM sodium pyruvate and 25 µg/mL kanamycin, supplemented with 10% FBS (growth medium) to final concentrations of 100, 20, 4 and 0.8 µM (two 24-well replica plates each). After BHK-21 cells were grown to confluency in 96-well plates, growth medium was replaced with fresh maintenance medium (growth medium with 1% inactivated FBS in place of 10% FBS) containing serially diluted test compound and HCoV-229E at a multiplicity of infection (MOI) of 0.01. Uninfected cells in the presence of serially diluted compound were used to assess the cytotoxicity of compounds. After a 3-day incubation at 37°C in a humidified 5% CO_2_ atmosphere, cell viability was determined by the MTT method (Pauwels et al., 1988). The effective concentration of test compound required to prevent virus-induced cytopathic effect (CPE) by 50% (EC_50_) and to cause 50% cell death in the absence of virus (CC_50_) were calculated by regression analysis.

### Antiviral assays in Huh-7 and RD cells

The antiviral activities of test compounds were evaluated against human coronaviruses alpha (229E), beta (OC43), MERS (EMC) and SARS (Urbani) using a neutral red assay to determine inhibition of virus-induced and compound-induced CPE and using a virus yield reduction (VYR) assay as a second, independent determination of the inhibition of viral replication.

#### Neutral red assay

Test compounds were dissolved in DMSO at a concentration of 10 mg/mL and serially diluted using eight half-log dilutions in test medium (Minimum Essential Medium supplemented with 5% FBS and 50 µg/mL gentamicin) so that the highest test concentration was 50 µg/mL. Each dilution was added to 5 wells of a 96-well plate with 80-100% confluent Huh-7 or RD cells (OC43 only). Three wells of each dilution were infected with virus, and two wells remained uninfected as toxicity controls. Six untreated wells were infected as virus controls and six untreated wells were left uninfected to use as virus controls. Viruses were diluted to achieve MOIs of 0.003, 0.002, 0.001 and 0.03 CCID_50_ per cell for 229E, OC43, MERS and SARS, respectively. Plates were incubated at 37±2°C in a humidified atmosphere containing 5% CO_2_.

On day 5 (229E and OC43) or day 7 (MERS and SARS) post-infection, when untreated virus control wells reached maximum CPE, the plates were stained with neutral red dye for approximately 2 hours (±15 minutes). Supernatant dye was removed, wells were rinsed with PBS, and the incorporated dye was extracted in 50:50 Sorensen citrate buffer/ethanol for >30 minutes and the optical density was read on a spectrophotometer at 540 nm. Optical densities were converted to percent of controls and the concentrations of test compound required to prevent virus-induced CPE by 50% (EC_50_) and to cause 50% cell death in the absence of virus (CC_50_) were calculated.

#### Virus yield reduction assay

Vero 76 cells were seeded in 96-well plates and grown overnight (37°C) to confluence. A sample of the supernatant fluid from each compound concentration was collected on day 3 post infection (3 wells pooled) and tested for virus titer using a standard endpoint dilution CCID_50_ assay and titer calculations using the Reed-Muench equation (1938) and the concentration of compound required to reduce virus yield by 90% (EC_90_) was determined by regression analysis.

### Antiviral assays in HAE cell preparations

The antiviral activities of test compounds were evaluated against SARS-CoV-2 (USA-WA1/2020) using made to order differentiated normal human bronchial epithelial (HAE) cells.

#### Cell Culture

HAE cells were grown on 6mm mesh disks and arrived in kits with either 12- or 24-well transwell inserts. During transportation the tissues were stabilized on a sheet of agarose, which was removed upon receipt. One insert was estimated to consist of approximately 1.2 × 10^6^ cells. Kits of cell inserts (EpiAirway™ AIR-100 or AIR-112) originated from a single donor, # 9831, a 23-year old, healthy, non-smoking, Caucasian male. The cells form polarized monolayers, the apical side of which is exposed to air and creates a mucin layer. Upon arrival, the cell transwell inserts were immediately transferred to individual wells of a 6-well plate according to the manufacturer’s instructions, and 1 mL of MatTek’s proprietary culture medium (AIR-100-MM) was added to the basolateral side, whereas the apical side was exposed to a humidified 5% CO_2_ environment. Cells were cultured at 37°C in a humidified atmosphere containing 5% CO_2_ for one day before the start of the experiment. After the 24-h equilibration period, the mucin layer, secreted from the apical side of the cells, was removed by washing with 400 µL pre-warmed 30 mM HEPES buffered saline solution 3X. Culture medium was replenished following the wash steps.

#### Viruses

Virus was diluted in AIR-100-MM medium before infection to yield a MOI when added to cultures of approximately 0.0015 CCID_50_ per cell.

#### Experimental design

Each compound treatment (120 μL) and virus (120 μL) was applied to the apical side, and the compound treatment (1 mL) was applied to the basal side. As a virus control, some of the cells were treated with cell culture medium only. After a 2-h infection incubation, the apical medium was removed, and the basal medium was replaced with fresh compound or medium (1 mL). The cells were maintained at the air-liquid interface. On day 5, cytotoxicity (CC_50_ values) in the uninfected, compound-treated inserts was estimated by visual inspection, and the basal medium was removed from all inserts and discarded. Virus released into the apical compartment of the HAE cells was harvested by the addition of 400 µL of culture medium that was pre-warmed at 37°C. The contents were incubated for 30 min, mixed well, collected, thoroughly vortexed and plated on Vero 76 cells for VYR titration. Separate wells were used for virus control and duplicate wells were used for untreated cell controls. Virus titers from each treated culture were determined as described above.

